# Distributed illusory figure-ground segmentation signatures across the dorsal and ventral streams

**DOI:** 10.1101/2023.11.01.565170

**Authors:** Ana Arsenovic, Anja Ischebeck, Natalia Zaretskaya

**Affiliations:** Department of Psychology, University of Graz, Graz, Austria; BioTechMed-Graz, Graz, Austria

**Keywords:** figure-ground segmentation, fMRI, illusory surface, topographic maps, dorsal stream, ventral stream

## Abstract

Visual illusions give rise to percepts that are more complex than the incoming sensory information, thereby exposing the inferential nature of perception. Early visual cortex (EVC) can represent not only physical input, but also illusory content of perception. Specifically, much like in real shape perception, during perception of an illusory shape the topographic representation of the shape’s surface is enhanced, while the regions surrounding it are suppressed, reflecting the figure-ground segmentation mechanisms on the neural level. It remains unclear whether such topographically specific figure-ground segmentation signatures are present in higher topographic maps along the visual hierarchy. To test this, we measured brain activity of 30 healthy human participants using functional magnetic resonance imaging (fMRI). The participants performed a central fixation task while passively observing a surrounding configuration of four inducers, which either formed or did not form an illusory shape. Using an additional functional localizer scan we could identify voxels uniquely representing the illusory surface and the background in the EVC, posterior parietal topographic region IPS0/V7 and several other maps along the dorsal and ventral streams. Apart from the EVC, we found signatures of figure-ground segmentation in several topographic maps. Illusory surface enhancement was observed in IPS0/V7, but without background suppression. Ventral topographic maps VO1 and VO2 showed background suppression only. Finally, other extrastriate areas, V3a, V3b and hV4, displayed both surface enhancement and background suppression. Our results demonstrate that topographically specific illusory shape responses are distributed along multiple topographic maps beyond the EVC. Furthermore, they point to the difference in neural signatures of figure-ground segmentation at various hierarchical levels and along different processing streams, stressing the independence of the surface enhancement and background suppression mechanisms.

## 1 Introduction

Our perceptual experience of the external world can be described as an ongoing inferential process. We receive the fragmented visual input from our retinas and transform it to produce a coherent visual percept, correctly segregating figures from the background. A great example of this inferential process are visual illusions such as Kanizsa figures (Kanizsa, 1976), in which the configuration of local pac-man elements gives rise to illusory contours that combine into illusory shapes.

Research has shown that the neural signatures of figure-ground segmentation for illusory shapes are quite similar to those of real shapes. Single cells in V1/V2 are well-known to respond to illusory contours in a way similar to the real contours (Maertens et al., 2008; von der Heydt et al., 1984; Zhou et al., 2000). The initial contour response is followed by a filling-in of a surface inside the figure and by a suppression of the background (Poort et al., 2016; Self et al., 2019). This pattern of surface enhancement and background suppression in response to illusory shapes has also been observed in human fMRI studies (Grassi et al., 2017; Kok & de Lange, 2014; Stoll et al., 2020). Both animal physiology and human fMRI findings consistently point to the role of feedback from higher-level areas in generating these responses (Kirchberger et al., 2021, 2023; Kok et al., 2016; Pak et al., 2020; Poort et al., 2016).

Indeed, neural correlates of shape perception are abundant in higher visual areas as well. Activity in the shape-representing areas such as the lateral occipital cortex (LOC) is typically enhanced when participants perceive a shape (Fang et al., 2008; Murray, 2004; Stanley & Rubin, 2003; Stoll et al., 2020). Accordingly, cortical feedback from the LOC has been shown to modulate illusory Kanizsa responses in the EVC (Chen et al., 2021; Wokke et al., 2013). Hence, signals from the shape-selective ventral stream regions are considered to be the source of feedback modulation in the EVC.

However, responses to illusory shapes at higher levels are present not only in the ventral stream regions, but also in areas of the dorsal processing stream. Specifically, the superior parietal cortex has been implicated in perceptual grouping and illusory shape perception based on dynamic and static visual cues (Arsenovic et al., 2022; Grassi et al., 2018; Liu et al., 2017; Robert et al., 2023; Zaretskaya et al., 2013). It has also been shown to represent the relations of individual object parts in an orientation-invariant manner (Ayzenberg & Behrmann, 2022b) and to influence the object representations in the ventral stream (Ayzenberg et al., 2023). There is thus substantial evidence that the dorsal stream plays an important role in perceptual grouping and object representation. It has been suggested that the dorsal stream is responsible for computing global shape representation, which is then transmitted to the ventral stream (Ayzenberg & Behrmann, 2022a).

But how exactly is global shape information represented in the dorsal stream? The human intraparietal sulcus consists of visual topographic maps, each representing one half of the visual field (Konen & Kastner, 2008a; Sereno et al., 2001; Silver & Kastner, 2009). It remains unclear how the shape representations within the IPS map onto the topographic representation of the visual space and whether a similar pattern of surface enhancement and background suppression can be found in the IPS maps as well. In the current study we examined the fine-scale activity within the topographic maps along the dorsal stream, including the IPS, and of the ventral stream, testing for neural signatures of figure-ground segmentation during the perception of illusory Kanizsa shapes and comparing the response pattern to the known activity pattern in V1. To test whether there is an interaction between the illusory shape representations in the early visual cortex and higher-level maps, we also examined the functional connectivity between the surface and background representations of V1 and the rest of the brain.

## 2 Materials and Methods

### 2.1 Participants

In total, 30 participants (12 male, 18 female) took part in the experiment (M = 23.53, SD = 2.92), which were part of our previous study (Arsenovic et al., 2022). Research questions of this and the previous study were jointly preregistered as one project on AsPredicted (https://aspredicted.org/ke4b9.pdf).

### 2.2 Stimuli

Visual stimuli were created with MATLAB R2019b (MathWorks, 2019) and Psychophysics Toolbox 3 (Brainard, 1997; Kleiner et al., 2007; Pelli, 1997) on Ubuntu 18.04 LTS Linux computer (for a detailed description of the experimental setup and the main experiment, see Arsenovic et al. (2022).

#### 2.2.1 Main Experiment

In the main experiment, participants were presented with four stimulus configurations (“no illusion,” “diamond,” “left triangle,” and “right triangle”, **Figure 1A**) for 12 seconds each (see **Figure 1B** for visualization of the example block). All stimulus configurations consisted of four pac-man inducers oriented such that they produced either an illusory diamond in the center, an illusory triangle on the left or on the right side, or no illusory figure. The order of configurations was pseudo-randomized. A central detection task (stream of blue and orange symbols) was present at all times to ensure central gaze fixation, as well as equal attention distribution across conditions. The central detection task was set to one of the two difficulty levels, easy or hard, throughout each run. In the easy task, participants were instructed to press a button on the response box in their hand each time the symbol color changed to blue (5% of all symbols). In the hard task, participants were instructed to press a button each time the plus symbol (“+”) appeared on the screen (5% of all symbols).

**Figure 1.**
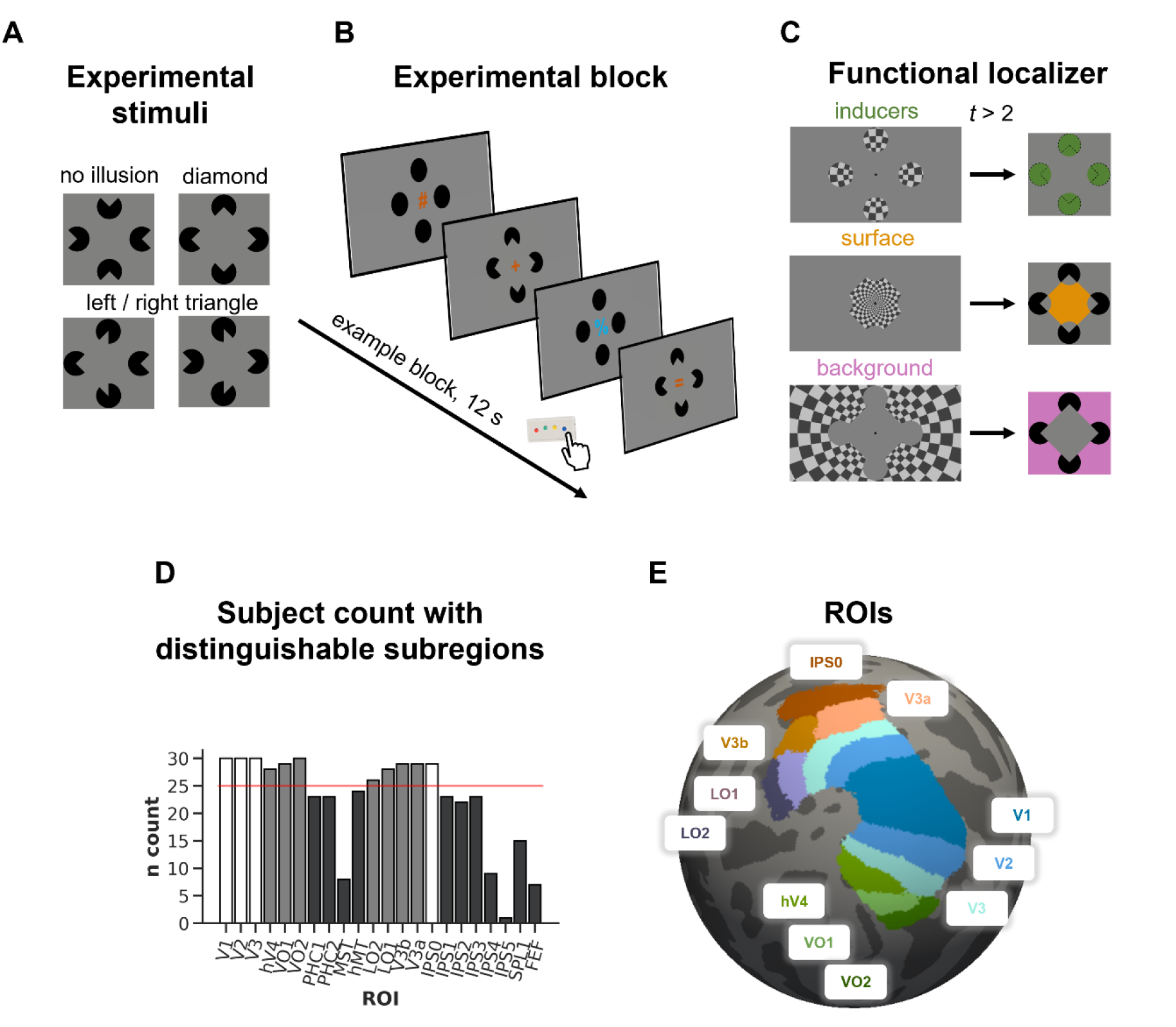
**A:** Stimulus conditions of the main experiment; **B:** Example block sequence in the main experiment, illustrated for easy detection task (detect blue symbols); **C:** Stimulus conditions of the functional localizer experiment (inducers, surface, background); **D:** count of subjects that have both surface and background localizer voxels in at least one hemisphere across topographic ROIs. Red line represents minimum sample size (25) calculated in the power analysis. White bars represent ROIs of the main analysis. Gray bars represent additional ROIs that also cross the power threshold and are included in the exploratory analysis. Black bars represent all other ROIs. **E:** Location of topographic maps that were tested for figure-ground modulation effects shown on left fsaverage spherical surface.

#### 2.2.2 Functional Localizer

After the main experiment, we acquired an additional functional localizer run for each participant. Three conditions were presented in the localizer run. Each condition consisted of a contrast-reversing (6 Hz) black-and-white checkerboard at 50% contrast presented as either circles at the location of the inducers (“inducers”), at the location of the emerging illusory shape in the center (“surface”) or at the location belonging neither to the inducers nor the illusory shape (“background”). Each of the four localizer circles was positioned at an eccentricity of 5.77° of the visual field and subtended 4.08°. The centrally positioned surface stimulus subtended 7.46° of visual angle (until the edges of the inducers). The background stimulus was presented outside of both surface and inducer stimuli, at a maximum eccentricity of 13.86° horizontally and 7.9° vertically (see **Figure 1C**).

Each localizer condition was presented 6 times for 12 seconds in a pseudorandomized and counterbalanced order (Brooks, 2012). The color detection task (easy task) was presented in the center of the screen to ensure fixation. In total, the localizer run lasted 216 seconds.

### 2.3 MRI data acquisition

All neuroimaging and behavioral data were acquired as part of a project with several research questions, part of which has been published (Arsenovic et al., 2022). A 3T Siemens MAGNETOM Vida MR scanner (Siemens Healthineers, Erlangen, Germany) with a 64-channel head coil was used to acquire all structural and functional scans. Functional MRI scans (both for the main experiment and the functional localizer) were acquired using the simultaneous multi-slice (SMS) accelerated echo-planar imaging (EPI) sequence using T2*-weighted blood-oxygenation-level-dependent (BOLD) contrast (58 axial slices, TR = 2000 ms, TE = 30 ms, FOV = 220 mm, flip angle = 82°, voxel size = 2.0 × 2.0 × 2.0 mm, SMS factor 2). To compensate for the EPI image distortion, we also acquired one EPI image with opposite phase encoding direction (P >> A). A high-resolution structural scan using the T1-weighted (T1w) MPRAGE sequence (TR = 2530 ms, TE = 3.88 ms, TI = 1200 ms, voxel size: 1.0 × 1.0 × 1.0 mm, GRAPPA factor 2) was also acquired.

### 2.4 MRI data preprocessing

Before further preprocessing of T1w and functional MRI data, we removed participants’ facial identifiers using pydeface 2.0.0 (Gulban et al., 2019). Data quality was ensured by visual inspection of individual and group reports via MRIQC 0.16.1 (Esteban et al., 2017). Structural and functional MRI data were preprocessed using fMRIPrep 20.2.3 (Esteban et al., 2019), which is based on Nipype 1.6.1 (Gorgolewski et al., 2011). We summarize the main preprocessing steps below.

#### 2.4.1 Structural MRI

The T1w image was corrected for intensity non-uniformity and used as T1w-reference, which was then skull-stripped. Cerebrospinal fluid (CSF), white-matter (WM) and grey-matter (GM) brain tissue segmentation was performed on the brain-extracted T1w image using FSL *fast* function (Woolrich et al., 2009). Additionally, volume-based spatial normalization to one standard space was performed using the T1w reference and ICBM 152 Nonlinear Asymmetrical template version 2009c (MNI152NLin2009cAsym). The latter was subsequently used for seed-based functional connectivity. Cortical surfaces were reconstructed using FreeSurfer *recon-all* stream (Dale et al., 1999).

#### 2.4.2 Functional MRI

A reference volume and its skull-stripped version were generated using a custom methodology of fMRIPrep. For each of the 9 BOLD runs (8 for the main experiment and 1 for the functional localizer) the following transformations were performed: A B0-nonuniformity map was estimated based on the EPI with opposite phase-encoding direction using AFNI’s *3dQwarp* (Cox, 1996). Based on the estimated susceptibility distortion, a corrected EPI reference was calculated for more accurate co-registration with the anatomical reference. The BOLD reference image was co-registered to the T1w reference image using FreeSurfer’s *bbregister* (Greve and Fischl, 2009). Additionally, head-motion parameters with respect to the BOLD reference image were estimated and slice-time correction was applied. These transforms were concatenated and applied in one interpolation step. The preprocessed BOLD time-series were resampled onto the individual’s anatomical reference generated from the T1w images. The preprocessed BOLD time-series were additionally resampled onto the MNI template (MNI152NLin2009cAsym).

### 2.5 Data analysis

#### 2.5.1 First-level GLM analysis

We performed volumetric general linear model (GLM) analyses using FreeSurfer’s FsFast. Because of a technical issue with the masks created in this version of fMRIPrep, we created custom functional masks by skull-stripping and then binarizing the template volume of each session (which corresponds to the first volume of the first run). All first-level analyses were performed on unsmoothed preprocessed BOLD runs. In addition to regressors of interest, we included run-specific effects and slow signal drifts as nuisance regressors.

The analysis of the main experiment was performed using the general linear model (GLM) approach. To account for potential differences in the shape of the hemodynamic response function (HRF) across areas, the latter was modelled as a finite impulse response (FIR) function with 6 time points (regressors) per condition. This approach allowed us to quantify BOLD responses within each ROI without assuming a specific response shape. Specifically, it allowed us to compare responses of the EVC and IPS areas, with the former showing a sustained and the latter a transient response to illusory shapes (Arsenovic et al., 2022). The GLM included eight conditions of interest (6 time point regressors each): 1) no illusion, 2) diamond, 3) left triangle, 4) right triangle during the hard task and conditions 5-8, which were the same stimulus conditions presented during the easy task. The illusory response at every time point was quantified by calculating six contrast estimates (CES) defined as the difference in beta estimate for each illusory shape condition (diamond, left triangle, and right triangle) and the “no illusion” condition, within the same task type. Since the task-specific activity is cancelled out in each contrast, the resulting CES represent the illusory shape response at each timepoint. The main statistical analysis was performed using the summed CES over all time points.

#### 2.5.2 Functional localizer GLM

A standard GLM was fit to the localizer data, with each of the three subregion conditions modelled as unique regressors (inducer, surface, background). All conditions were modelled as full-duration 12 s blocks and were convolved with a canonical HRF. The resulting beta estimates were then used in a custom MATLAB script to extract voxels that respond uniquely to each of the subregion conditions (see Statistical analysis).

#### 2.5.3 Regions-of-interest (ROI) definition

All regions of interest were defined in each participant’s individual anatomical space. First, maximum probability maps (MPM) of the surface-based probabilistic atlas (Wang et al., 2015) in fsaverage space (Benson & Winawer, 2018) were resampled into individual surface space using *mri_surf2surf*. Then, each resulting surface output was resampled to volume with *mri_surf2vol* via the method of filling the cortical ribbon using ribbon.mgz. Finally, each volumetric output was resampled into individual (T1w-aligned) template functional volume space using *mri_label2vol*.

The results of the localizer GLM were used to define voxels representing each subregion within each visual topographic map. Specifically, we used t-statistic maps for the contrast comparing each condition’s activity with the average of the remaining two conditions. Voxels exceeding a t-value of 2 were retained. These voxels were used to average activity within each subregion during the main experiment. Participants were excluded from the analysis if no voxels for the surface and the background subregion in at least one hemisphere could be identified (the presence of voxels in the inducer subregions was not considered due to their relatively small size).

For the bilateral presentation, we averaged the mean activity from both hemispheres. For the contralateral presentation, we averaged responses from the right hemisphere during left triangle presentation and responses from the left hemisphere during right triangle presentation. Finally, for the ipsilateral presentation, we averaged the responses from the right hemisphere during right triangle presentation and responses from the left hemisphere during left triangle presentation. In all three presentations, if subregion voxels were present only in one hemisphere, the mean activity from that hemisphere was used.

#### 2.5.4 Statistical analysis

Statistical analyses were performed in RStudio version 4.1.2 (RStudio Team, 2021). First, we determined which of the topographic maps are suitable for the classical statistical inference by performing power analysis using GPower 3.1.9.7 (Faul et al., 2007). We determined the minimum number of participants needed to reach at least 80% statistical power for a one-tailed paired samples t-test, with alpha 0.05, expected effect size of 0.59. The expected effect size was determined from previously reported effect size within the posterior parietal cortex related to responses to illusory percepts (Grassi et al., 2018). This resulted in a minimum *n* = 25 of participants with at least one voxel activated in both surface and background (see **Figure *1*D**), leading to inclusion of areas V1, V2, V3 and IPS0 in the main statistical analysis. For completeness, we also include other topographic ROIs that had 25 or more participants in the results (hV4, VO1, VO2, LO1, LO2, V3a, V3b, **Figure 1E**). Additionally, all IPS topographic maps (IPS0-IPS4) were analyzed using an equivalent Bayesian approach, to account for the reduction of statistical power in the parietal maps above IPS0.

##### 2.5.4.1 Classical inferential statistics

To test for subregion-specific illusory responses, we performed 2 × 2 repeated-measures ANOVA within each ROI, with factors “subregion” (surface, background) and “task” (easy, hard), separately for the bilateral, contralateral, and ipsilateral stimulus conditions.

##### 2.5.4.2 Bayesian estimation

To quantify the relative predictive performance of the null and alternative hypotheses in IPS0-IPS4, we calculated Bayes factor (Wagenmakers, 2007) using BayesFactor R package version 0.9.12-4.4 (Morey et al., 2022), which represents the weight of evidence for observing the experimental data under the null and under the alternative hypotheses. The descriptions of cutoffs for evidence in the results were done according to Wagenmakers et al. (2018).

We computed all model estimates within each ROI at once using *anovaBF* with one million Monte Carlo simulations each. We modelled the participants as an additive effect on top of the other effects (as is typically assumed). To determine if there was a difference in response across subregions under different task difficulties, we specified a 2 × 2 Bayesian ANOVA, with factors “subregion” and “task”. We report the performances of all tested models (subregion, task, subregion + task, and subregion + task + interaction) for every topographic map. For each model, we report BF_10_ – the Bayes factor representing evidence for H_1_ (existence of effect) over H_0_ (no effect).

#### 2.5.5 Functional connectivity

Functional connectivity (FC) analysis was performed in CONN toolbox (Whitfield-Gabrieli & Nieto-Castanon, 2012) version 21.a (Nieto-Castanon & Whitfield-Gabrieli, 2021) and SPM12 (Penny et al., 2011). First, preprocessed fMRI data from the main experiment were imported into CONN, along with six fMRIPrep-generated realignment parameters. The only additional preprocessing step done in CONN was outlier detection using default settings. Outlier volumes along with the four initial dummy volumes were entered as covariates in the first-level GLM.

MNI-space functional data were smoothed using spatial convolution with a Gaussian kernel of 8 mm full width half maximum (FWHM). In addition, functional data were denoised using a standard denoising pipeline (Nieto-Castanon, 2020) including the regression of potential confounding effects characterized by CSF timeseries (5 CompCor noise components), motion parameters and their first order derivatives (12 regressors, Friston et al., 1996), outlier scans (maximum 28, Power et al., 2014), white matter timeseries (16 CompCor noise components), as well as run-specific offset, linear and quadratic trends (3 regressors) within each functional run, followed by high-pass frequency filtering of the BOLD timeseries (Hallquist et al., 2013) above 0.008 Hz. CompCor (Behzadi et al., 2007; Chai et al., 2012) noise components within white matter and CSF were estimated by computing the average BOLD signal as well as the largest principal components orthogonal to the BOLD average, motion parameters, and outlier scans within each subject’s eroded segmentation masks.

##### 2.5.5.1 ROI-to-ROI functional connectivity

Following our primary hypothesis on the involvement of the dorsal stream regions in producing figure-ground modulation in the EVC, we first conducted a ROI-to-ROI connectivity (RRC) analysis, where we estimated FC matrices between each pair of the eight subregions of interest (merged left and right hemisphere ROI in V1, V2, V3, and IPS0, with surface and background subregion in each). Subregion time courses were represented by an average time course of all voxels in subregion in individual T1-space unsmoothed data. FC strength was represented by Fisher-transformed bivariate correlation coefficients between residuals of a weighted GLM (Nieto-Castanon, 2020) for each ROI pair after accounting for condition-related activity. Condition-related activity was modelled out from the ROI’s time course by recreating our FIR GLM setup of the main fMRI analysis. Connectivity values of individual time points were summed, yielding one value per subject, per connection and per condition. Group-level inferences were performed using a paired t-test between illusory and inverted conditions at each individual functional connection. The FDR-corrected connection-level threshold at p < 0.05 was applied to the results (Benjamini & Hochberg, 1995).

##### 2.5.5.2 Seed-based functional connectivity

In an additional exploratory analysis of potential figure-ground modulation sources over the entire brain, we conducted a seed-based functional connectivity analysis (SBC). Seed regions included V1 surface and V1 background within each hemisphere (derived as an average time course of all voxels in subregion in individual T1-space unsmoothed data). Functional connectivity strength was represented by Fisher-transformed bivariate correlation coefficients between residuals of a weighted GLM (Nieto-Castanon, 2020). GLM was estimated separately for each seed area and target voxel, modeling the association between their BOLD signal timeseries. Condition-related activity was modelled out from the voxel’s time course by recreating our FIR GLM setup from the ROI analysis. Group-level analyses were performed using second-level GLM (Nieto-Castanon, 2020). GLM was estimated for each voxel, with first-level connectivity measures (one measurement per subject, condition and time point) as dependent variables. Voxel-level hypotheses were evaluated using a t contrast comparing each shape condition with the inverted condition, summing values of all time points. Cluster-level inferences were based on parametric statistics from the Gaussian Random Field theory (Nieto-Castanon, 2020; Worsley et al., 1996). We applied a threshold on the results using a combination of a cluster-forming threshold at p < 0.001, and FDR-corrected cluster-size threshold at p < 0.05 (Chumbley et al., 2010). We computed SBC for bilateral > no illusion, contralateral > no illusion and ipsilateral > no illusion contrasts of the easy task within each hemisphere, for each V1 subregion seed. Because IPS0 showed up as the only intraparietal topographic map with illusory surface responses (see Results), in an additional analysis we also computed SBC with surface and background subregions of IPS0 in each hemisphere.

## 3 Results

### 3.1 Topographic illusory responses in IPS0 and early visual areas

We tested whether IPS0 shows a topographically specific response to illusory shapes, comparing the response pattern to the well-known response pattern in the early visual cortex (Grassi et al., 2017; Kok & de Lange, 2014). Consistent with previous findings, we observed a significant difference between responses in the illusory surface subregion and in the background subregion, with the surface subregion showing a higher response compared to the background. This effect was present for all illusory shape presentations in V1, V2, as well as in V3 (all p-values < 0.001, ***Supplementary Table 1***). No topographic map of the EVC showed an effect of the task or an interaction between the task and the subregion.

We further examined whether the above effects were driven by illusory surface enhancement, background suppression, or both. Since there was no effect of task, we considered responses for the easy task only. This analysis revealed that in all three topographic maps of the EVC, the representation of the illusory surface was enhanced, and that of the background suppressed for the bilateral and the contralateral shape presentations. For the ipsilateral presentation, however, only the background suppression was responsible for the differences between subregions (**Figure 2**, ***Supplementary Table 2***).

**Figure 2.**
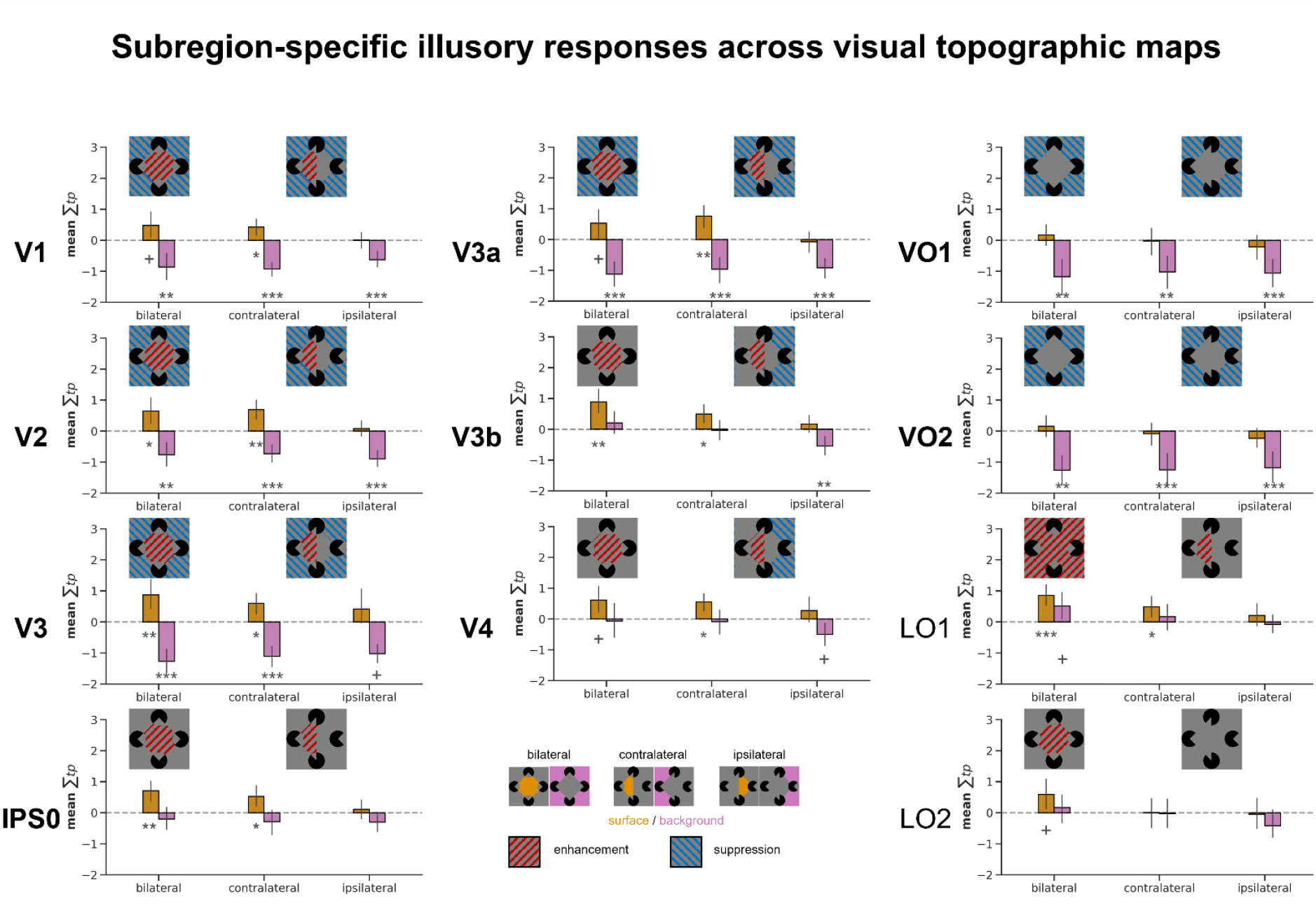
Subregion-specific illusory shape responses across visual topographic maps. Left column: Illusory responses in early visual areas and the IPS0; Middle and right column: Illusory responses across topographic maps in other analyzed visual areas. Areas with a significant figure-ground modulation effect (see also **Table 1**) are shown in **bold** font. Asterisks indicate the significance of a one-sample t-test for a nonzero illusory shape response in each subregion: ***p<0.001, **p<0.01, *p<0.05, Bonferroni-Holm corrected for 11 t-tests, +p<0.05 uncorrected. Error bars represent 95% CIs. Color of the bars (surface – orange and background – purple) represent subregions of each stimulus presentation. Striped filling of visualized figures represents significant enhancement (red) or suppression (blue) response to illusory vs. non-illusory presentation (uncorrected t-test) in that specific subregion.

The pattern of responses within IPS0 was similar, but not identical to early visual areas. Specifically, similarly to the EVC, IPS0 also showed a difference between subregions, with a higher response in the illusory surface compared to the background (bilateral: *F*_(1, 28)_ = 18.592, *p* < .001, *η_p_^2^* = 0.399; contralateral: *F*_(1, 28)_ = 11.013, *p* = .003, *η_p_^2^* = 0.282; ipsilateral: *F*_(1, 28)_ = 22.907, *p* < .001, *η_p_^2^* = 0.45). Like in the EVC, bilateral and contralateral, but not the ipsilateral presentation elicited an enhanced response in the subregion representing illusory surface. However, in contrast to early visual areas, we did not observe background suppression in any of the illusory shape presentations (**Figure 2**). There was no effect of the task and no interaction between the task and the subregion.

### 3.2 Illusory responses in other visual areas

As an exploratory analysis, we also investigated illusory responses within other topographic visual areas, where both surface and background subregions could be identified in at least 25 subjects (as calculated by the power analysis). These areas include the ventral and dorsal mid-level topographic maps hV4, VO1, VO2, LO1, LO2, V3a and V3b. Ventral topographic maps hV4, VO1 and VO2, as well as V3a and V3b showed response differences between subregions (**Table 1**). Furthermore, LO1 showed a marginally significant interaction between subregion and task in the bilateral presentation (*F*_(1, 27)_ = 4.24, *p* = 0.049, *η_p_^2^* = 0.136) and a task effect for the ipsilateral presentation (*F*_(1, 27)_ = 5.199, *p* = 0.031, *η_p_^2^* = 0.161) and LO2 showed no effects of either subregion or task (all *p*-values ≥ 0.145).

**Table 1.**
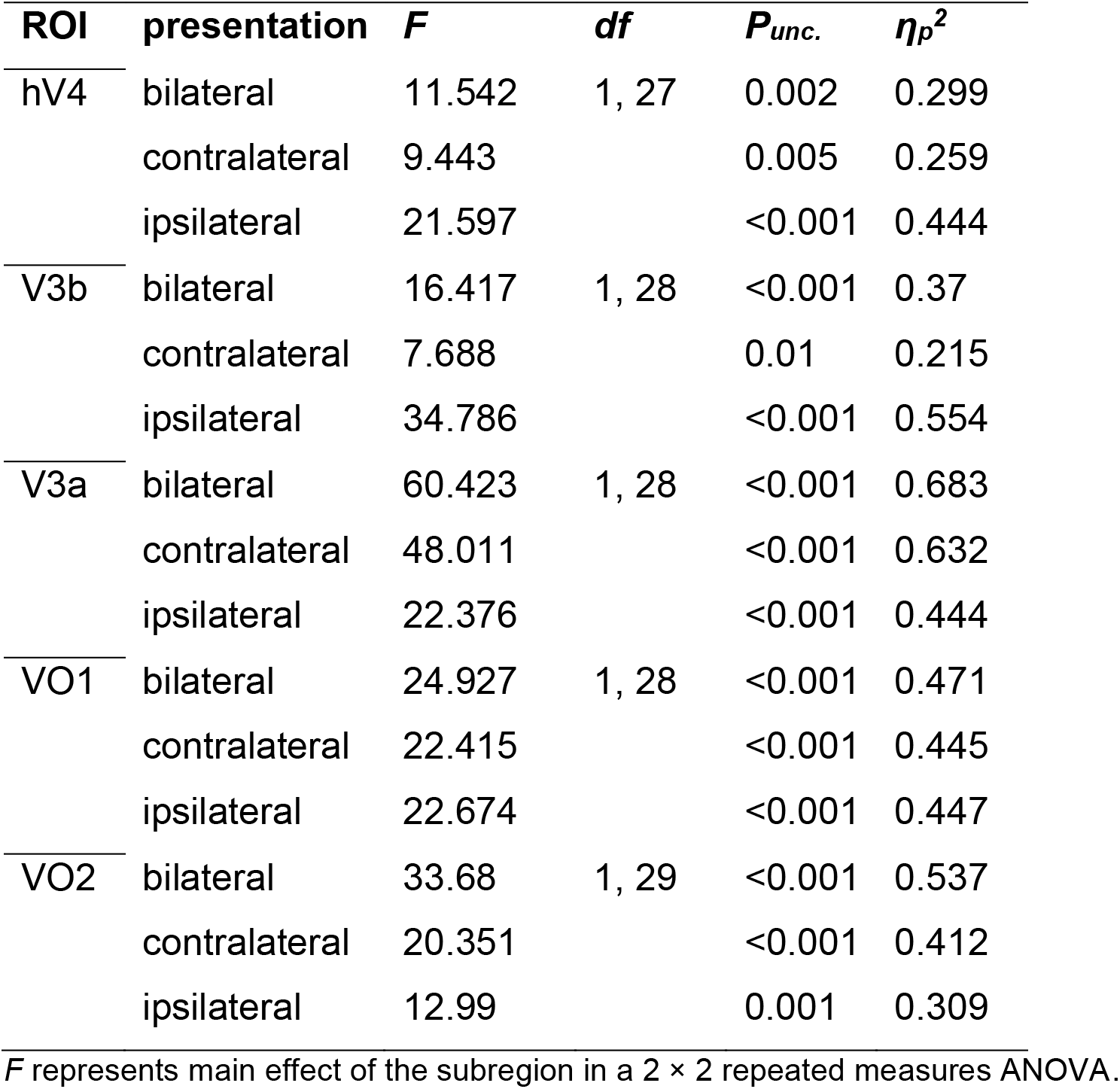
Figure-ground modulation signatures for different illusory shape presentations in other visual areas.

| ROI | presentation | *F* | *df* | *P<sub>unc.</sub>* | $\eta_p^2$ |
| --- | --- | --- | --- | --- | --- |
| hV4 | bilateral | 11.542 | 1, 27 | 0.002 | 0.299 |
|  | contralateral | 9.443 |  | 0.005 | 0.259 |
|  | ipsilateral | 21.597 |  | <0.001 | 0.444 |
| V3b | bilateral | 16.417 | 1, 28 | <0.001 | 0.37 |
|  | contralateral | 7.688 |  | 0.01 | 0.215 |
|  | ipsilateral | 34.786 |  | <0.001 | 0.554 |
| V3a | bilateral | 60.423 | 1, 28 | <0.001 | 0.683 |
|  | contralateral | 48.011 |  | <0.001 | 0.632 |
|  | ipsilateral | 22.376 |  | <0.001 | 0.444 |
| VO1 | bilateral | 24.927 | 1, 28 | <0.001 | 0.471 |
|  | contralateral | 22.415 |  | <0.001 | 0.445 |
|  | ipsilateral | 22.674 |  | <0.001 | 0.447 |
| VO2 | bilateral | 33.68 | 1, 29 | <0.001 | 0.537 |
|  | contralateral | 20.351 |  | <0.001 | 0.412 |
|  | ipsilateral | 12.99 |  | 0.001 | 0.309 |

*F* represents main effect of the subregion in a 2 × 2 repeated measures ANOVA.

To test whether the effects in other topographic maps were driven by surface enhancement, background suppression, or both, we again tested responses in each subregion against zero. Comparably to the main analysis, we considered the illusory shape responses in the easy task only. Neither VO1 nor VO2 showed illusory surface enhancement (all *p*-values > 0.171), and both topographic maps showed background suppression for all three stimulus presentations (all *p*-values < 0.002). On the other hand, similarly to the EVC, we found surface enhancement for the bilateral and contralateral presentation in hV4 (bilateral *p* = 0.011, contralateral *p* = 0.001), and a background suppression, but only for the ipsilateral presentation (*p* = 0.015). Ipsilateral surface and bilateral/contralateral background were not significantly modulated by the illusory response (all p-values > 0.222). V3a showed a surface enhancement for the bilateral (*p* = 0.0345) and contralateral presentations (*p* < 0.001) and background suppression in all shape presentations (all *p*-values < 0.001). Finally, V3b also showed surface enhancement in bilateral (*p* < 0.001) and contralateral presentations (*p* = 0.004), and a background suppression only in the ipsilateral presentation (*p* = 0.001). Responses of all analyzed topographic maps are shown in **Figure 2** for each shape presentation and subregion (easy task). In sum, figure-ground modulation effects in different maps seem to be driven by different processes. While IPS0 shows illusory surface enhancement, maps VO1 and VO2 show background suppression, while areas V3a, V3b, hV4 and the early visual areas V1-V3 show both types of responses.

### 3.3 Parietal topographic maps IPS0-IPS4

Additionally, to make sure we did not miss any potential effects due to the lack of statistical power, we tested all topographic interparietal areas by performing 2 × 2 repeated-measures Bayesian ANOVA, with participant as random (nuisance) factor and subregion and task as fixed factors. First, we confirmed the results obtained with classical statistical inference in IPS0. The most successful model for data in IPS0 is the effect of the subregion, where we note extreme (diamond, ipsilateral, BF_10_ > 100) or very strong (contralateral, BF_10 =_ 77.99) evidence for a difference between the surface and the background subregion for all three stimulus presentations. There is no evidence for an effect or an interaction of the two effects for data in IPS1, IPS2 or IPS3. On the other hand, the most successful performing model in IPS4 is the task effect model, which shows strong evidence (BF_10_ = 10.18) for differences in illusory response across task difficulty, despite the low number of participants included (see **Supplementary Tables 3, 4, 5, 6** and **7** for the results within each shape presentation for each calculated model in IPS0-4.). Performance of different models is presented in ***Figure 3*** bellow.

**Figure 3.**
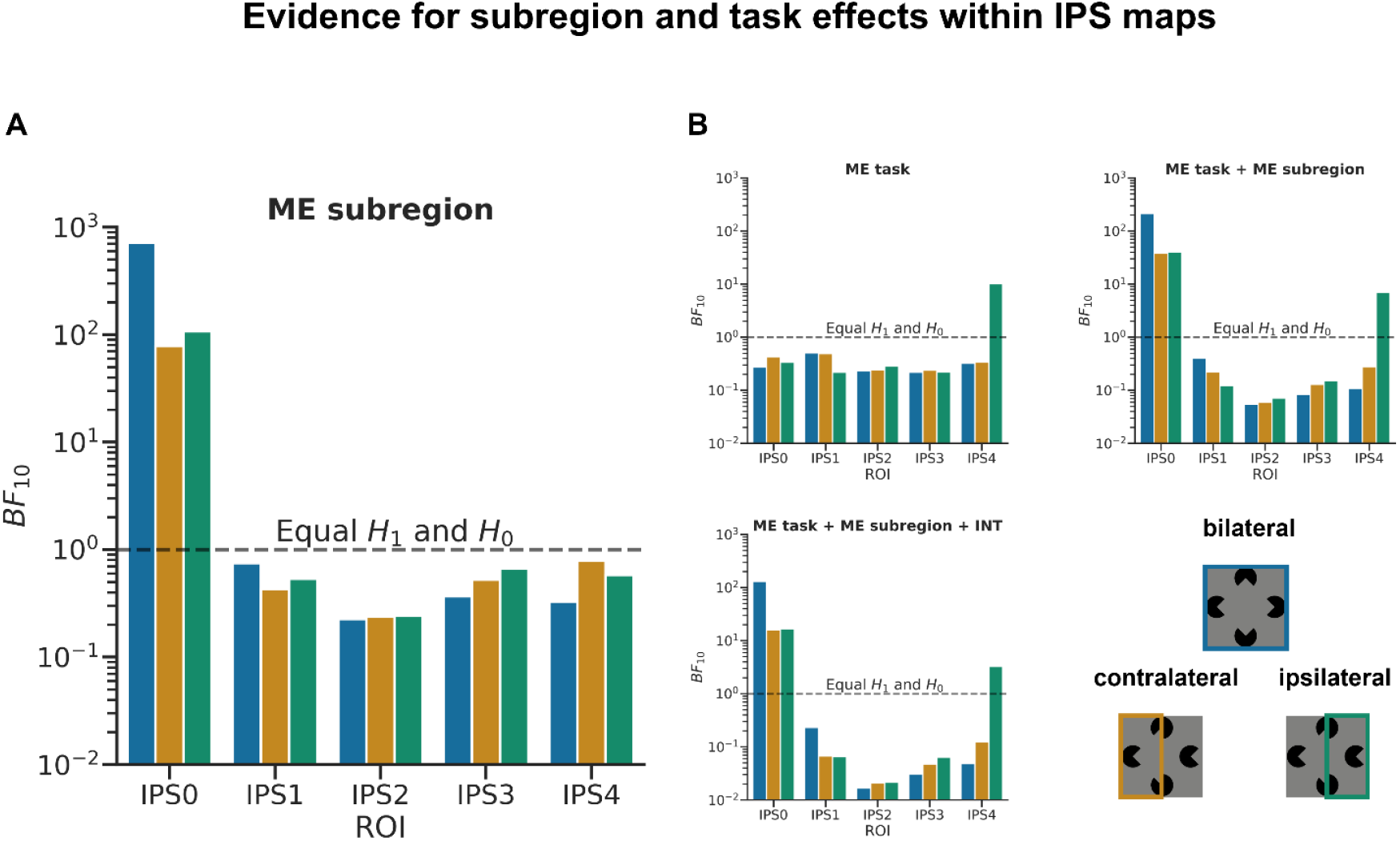
Results of the 2 × 2 Bayesian ANOVA for each area of the intraparietal sulcus (IPS0-4); ***A:*** Main effect of the subregion as the most successful model for data in IPS0 across all stimulus presentations; ***B:*** all other calculated models; ME: main effect, INT: interaction of main effects. *Colored bars represent each illusory shape presentation as shown in the bottom right (bilateral – blue, contralateral – orange, and ipsilateral – green)*.

### 3.4 Functional connectivity

To examine whether surface enhancement and background suppression across the visual hierarchy arise through an interaction between higher and lower-level topographic areas, we performed functional connectivity analyses. We calculated differences in functional connectivity between each illusory stimulus condition and the no illusion condition, while accounting for the stimulus-evoked activity using an FIR GLM model.

We found no significant connectivity changes in the diamond condition compared to no illusion condition in our ROI-to-ROI analysis that included V1, V2, V3 and IPS0 surface and background subregions of both hemispheres.

To examine whether surface enhancement and background suppression in topographic regions arise from an interaction with other, potentially non-topographic regions of the brain, we performed a seed-based functional connectivity analysis, relating fluctuations of activity in each subregion with those of each voxel in the brain in different conditions. The comparison of the left and right triangle vs. inverted yielded no significant connectivity differences, neither for illusory surface nor for the background seeds. However, we observed a significant connectivity reduction of both the left and the right V1 seeds representing illusory surface in the bilateral illusory surface condition (illusory diamond). Both left and the right V1 seeds reduced their functional connectivity with the same lateral parietal default-mode network region of the right hemisphere (DMN LP). For the right V1 seed, 191 voxels (90%) of the negative cluster (212 voxels, *p_FDR_* = 0.001) with a peak at MNI [+42 -64 +28] overlapped with the DMN LP in the right hemisphere (**Figure 4A**). For the left V1 seed, 118 voxels (91%) of the negative cluster (129 voxels, *p_FDR_* = 0.023) with a peak at MNI [+44 -64 +30], overlapped the same DMN LP in the right hemisphere (**Figure 4B**). We did not find significant changes in functional connectivity of the background seed in V1 in either hemisphere. Surface and background seeds in IPS0 yielded no significant effects either.

**Figure 4.**
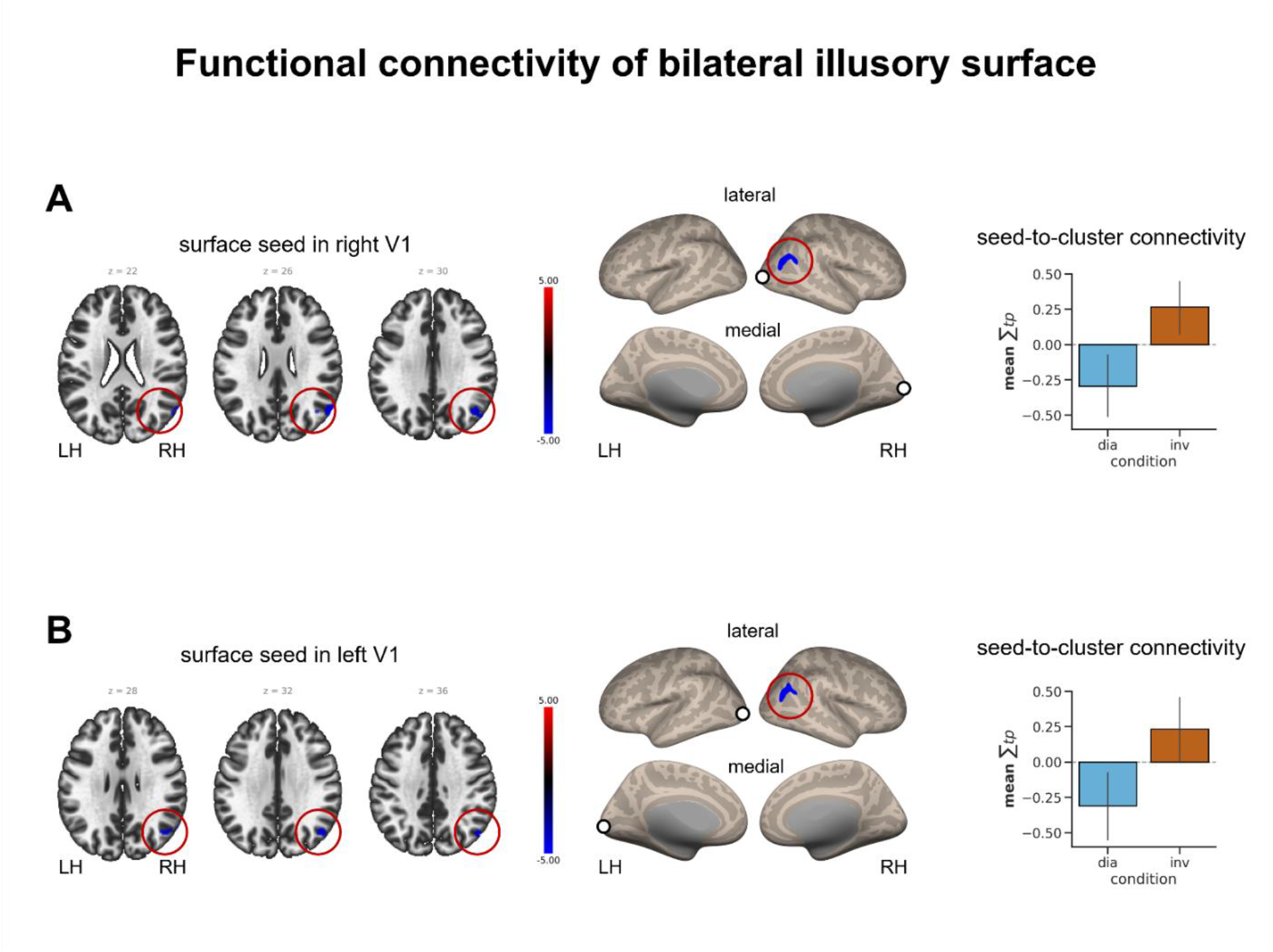
Results of the seed-based functional connectivity; **A** – seed in illusory surface subregion of the right V1; **B** – seed in illusory surface subregion of the left V1. Clusters are overlayed on the MNI template brain (left) and on the inflated surfaces (middle). Bar plots show connectivity values within the significant cluster (sum of all time points, right) LH: left, RH: right hemisphere.

## 4 Discussion

### 4.1 Summary

In this study, we performed a detailed investigation of the illusory shape representation across visual cortical topographic maps. We report a topographically specific activity pattern during illusory shape perception in several maps up to and including IPS0. Specifically, we show that along with the EVC, extrastriate maps hV4, V3a and V3b exhibit both surface enhancement and background suppression, VO1 and VO2 show background suppression only, and V7/IPS0 shows only surface enhancement. Our results indicate the presence of figure-ground segmentation effects at multiple levels across the visual hierarchy and emphasize the relative independence of the processes that drive figure enhancement and background suppression. Crucially, they point to the specific role of the dorsal stream, in particular IPS0, in representing illusory surface.

### 4.2 Dorsal stream contribution to illusory shape perception

Our results of the figure-ground segmentation effects up to the area IPS0 along the dorsal stream add to the growing body of evidence for the dorsal stream contribution to real (Ayzenberg & Behrmann, 2022a, 2022b; Freud et al., 2016, 2017; Konen & Kastner, 2008b), as well as illusory shape processing (Arsenovic et al., 2022; Grassi et al., 2018). One hypothesis about the exact contribution of the dorsal stream in general, and parietal cortex in particular, is that it provides a coarse skeletal-like shape representation that is stripped of local features (Ayzenberg & Behrmann, 2022a). Crucially, the dorsal stream is thought to encode the spatial arrangement of individual shape elements, but is largely invariant to the overall object orientation (Ayzenberg & Behrmann, 2022b). Our results in IPS0 and V3a are not consistent with this notion, since orientation-independence is incompatible with the topographic specificity we find in these areas. Furthermore, Pac-man inducers that produce the left and right triangles in our study essentially represent the same object in different orientations. An orientation-invariant representation would yield similar responses to the contralateral and ipsilateral shapes, which was not the case in these areas. It is possible, however, that V3a and IPS0 are too low along the dorsal stream hierarchy to reflect its characteristic features during shape processing. In our previous study, responses that were independent of the side of the illusory triangle were observed more anterior within the intraparietal sulcus, in the topographic maps IPS3 and IPS4 (Arsenovic et al., 2022). Hence, it is possible that the representation of objects becomes more and more coarse and orientation-invariant along the posterior-anterior gradient within the IPS.

What is the functional role of the topographic representation of illusory shapes in the dorsal stream? Electrophysiology recordings in macaques revealed that the topographic activity in the primary visual cortex emerges over several stages. After the initial contour enhancement, there is an enhancement of activity over the whole surface of the figure, which is followed by the background suppression (Poort et al., 2016; Self et al., 2019). Notably, both surface enhancement and background suppression are confined to the superficial/deep cortical layers, which are the primary targets of feedback inputs to V1 (Poort et al., 2016). Our results show that the IPS0 map exhibits figure enhancement only, while the ventral stream regions VO1 and VO2 exhibit only background suppression. The feedback signal in V1 may thus stem from different sources depending on what part of the scene is represented: first, the topographic maps of the dorsal stream may provide the feedback signal related to the extracted surface, while later the topographic maps of the ventral stream may provide signal related to the suppressed background. This idea is consistent with the faster and more automatic processing in the dorsal stream (Ayzenberg et al., 2023; Liu et al., 2017). The hypothesis about the differential feedback influence of dorsal and ventral stream in figure-ground segregation needs to be tested more directly in the future, perhaps using high-resolution fMRI acquisitions at different cortical depths (Jia et al., 2023; Zaretskaya, 2021). In any case, our study supports the previous observations that the surface enhancement and background suppression during figure-ground modulation is mediated by different mechanisms (Grassi et al., 2017; Poort et al., 2016; Strother et al., 2012).

### 4.3 Default-mode network involvement in shaping subjective visual experience

Interestingly, our exploratory functional connectivity analysis did not show a connectivity increase between V1 and other shape-processing areas. Instead, we found connectivity change between the representation of illusory surface and the right-hemispheric lateral-parietal node of the DMN. The correlation between these two areas changed from being positive or zero in the inverted condition to being below zero in the diamond condition (bar graphs in **Figure 4**). During illusory surface perception, activity in the surface representation of V1 and the DMN cluster thus fluctuated in opposite directions.

DMN activity is typically associated with tasks unrelated to sensory processing such as mind wandering and abstract thought (Fox et al., 2015). However, more recently, it has been repeatedly linked to shaping subjective aspects of perceptual content. For example, increased activity in the DMN nodes, including the lateral parietal cortices, has been observed during the disambiguation of degraded visual stimuli, and is thought to reflect the influence of prior experience on perception (González-García et al., 2018). Furthermore, a recent resting-state fMRI study has shown that the DMN connectivity to V1 predicted the dominant percept in binocular rivalry (Lyu et al., 2022). Finally, parietal connectivity with the DMN during resting-state was associated with an individual tendency to perceive the illusory shape interpretation of a global-local bistable stimulus (Wilding et al., 2023). What we see as functional connectivity between the illusory surface representation and one of the DMN nodes may therefore be a manifestation of the DMN’s contribution to perception.

A further core feature of the DMN is the presence of negative, but at the same time topographically specific visual responses, which were revealed using the modeling of voxel population receptive fields (pRFs, Klink et al., 2021; Szinte & Knapen, 2020). The functional role of these responses remains entirely unclear. It would be interesting to test whether the pRF position of voxels in the lateral parietal DMN node we identified corresponds to the topographic representation of the illusory surface. Such a finding would indicate a communication between the topographic representation of the illusory surface at the two ends of the cortical hierarchy, V1 at the lowest level and DMN at the highest level (Konishi et al., 2015; Margulies et al., 2016). An interesting hypothesis to test in futures studies is whether the DMN connectivity is unique to illusory shape processing or is also present during processing of real shapes. The former would confirm its contribution to shaping subjective aspects of perception.

In sum, our connectivity results point to the potential DMN involvement in illusory shape processing. Taking into account and understanding the contribution of the default-mode network to illusory aspects of perception is an interesting avenue for future investigation.

### 4.4 Conclusion

Our study demonstrates the presence of figure-ground segmentation effects during illusory shape perception in several visual topographic areas of the dorsal and ventral streams, including the intraparietal area IPS0. The illusory response pattern in IPS0 had both similarities and differences with the early visual areas. Similarly to the EVC, IPS0 showed an enhancement of the illusory shape representation, but differently from the EVC there was no background suppression, which was in contrast present in the ventral stream maps VO1 and VO2. A distributed and subregion-specific pattern of responses to illusory figures is thus present in multiple topographic maps along the dorsal and the ventral streams beyond the early visual cortex. Future studies using higher spatial resolution may be able to reveal topographically specific responses to illusory shapes in other, higher- and lower-level visually responsive areas beyond the ones described here. They should also focus on discerning the specific signatures of illusory, as opposed to real shape processing.

## Data and Code Availability

All analysis scripts and statistical data are available at https://osf.io/zq9pw/. BIDS compatible whole-brain fMRI dataset is available at https://osf.io/rvf27/.

## Author Contributions

A.A.: Conceptualization, Methodology, Investigation, Data curation, Software, Formal analysis, Visualization, Writing - original draft, and Writing - review & editing

A.I.: Writing - review & editing, Supervision

N.Z.: Conceptualization, Methodology, Investigation, Writing - review & editing, Funding acquisition, Supervision

## Acknowledgements

We thank Thomas Zussner for the help with MRI acquisition.

## Supplementary data

### Topographic illusory responses in IPS0 and early visual areas

**Supplementary Table 1.** Differences between surface and background responses to illusory shapes in the EVC.

| ROI | presentation | $F_{(1, 28)}$ | $p_{unc.}$ | $\eta_p^2$ |
| --- | --- | --- | --- | --- |
| V1 | bilateral | 132.679 | < .001 | 0.826 |
|  | contralateral | 129.819 | < .001 | 0.823 |
|  | ipsilateral | 48.436 | < .001 | 0.634 |
| V2 | bilateral | 97.07 | < .001 | 0.776 |
|  | contralateral | 109.219 | < .001 | 0.796 |
|  | ipsilateral | 82.701 | < .001 | 0.747 |
| V3 | bilateral | 137.453 | < .001 | 0.831 |
|  | contralateral | 124.096 | < .001 | 0.816 |
|  | ipsilateral | 82.428 | < .001 | 0.746 |

$F$ represents main effect of the subregion in a $2 \times 2$ repeated measures ANOVA

**Supplementary Table 2.** Illusory shape response in the subregions of the EVC and IPS0 (presentation > no illusion)

| ROI / subregion | presentation | $t_{(28)}$ | $p_{unc.}$ | Cohen's $d$ |
| --- | --- | --- | --- | --- |
| **V1** |  |  |  |  |
| surface | bilateral | 2.211 | 0.035 | 0.411 |
|  | contralateral | 3.101 | 0.004 | 0.576 |
|  | ipsilateral | 0.066 | 0.948 | 0.012 |
| background | bilateral | -4.26 | < 0.001 | -0.791 |
|  | contralateral | -7.734 | < 0.001 | -1.436 |
|  | ipsilateral | -5.018 | < 0.001 | -0.932 |
| **V2** |  |  |  |  |
| surface | bilateral | 3.088 | 0.005 | 0.573 |
|  | contralateral | 4.516 | < 0.001 | 0.839 |
|  | ipsilateral | 0.604 | 0.55 | 0.112 |
| background | bilateral | -3.944 | < 0.001 | -0.732 |
|  | contralateral | -5.097 | < 0.001 | -0.946 |
|  | ipsilateral | -7.023 | < 0.001 | -1.304 |
| **V3** |  |  |  |  |
| surface | bilateral | 3.599 | 0.001 | 0.668 |
|  | contralateral | 3.475 | 0.002 | 0.645 |
|  | ipsilateral | 1.388 | 0.176 | 0.258 |
| background | bilateral | -6.191 | < 0.001 | -1.15 |
|  | contralateral | -6.445 | < 0.001 | -1.197 |
|  | ipsilateral | -6.691 | < 0.001 | -1.242 |
| **IPS0** |  |  |  |  |
| surface | bilateral | 4.003 | < 0.001 | 0.743 |
|  | contralateral | 2.951 | 0.006 | 0.548 |
|  | ipsilateral | 0.698 | 0.491 | 0.13 |
| background | bilateral | -1.116 | 0.274 | -0.207 |
|  | contralateral | -1.378 | 0.18 | -0.256 |
|  | ipsilateral | -1.874 | 0.071 | -0.348 |

Two-tailed one sample *t*-test comparing mean CES against 0.

### Bayesian models performance IPS0-IPS4

**Supplementary Table 3.**
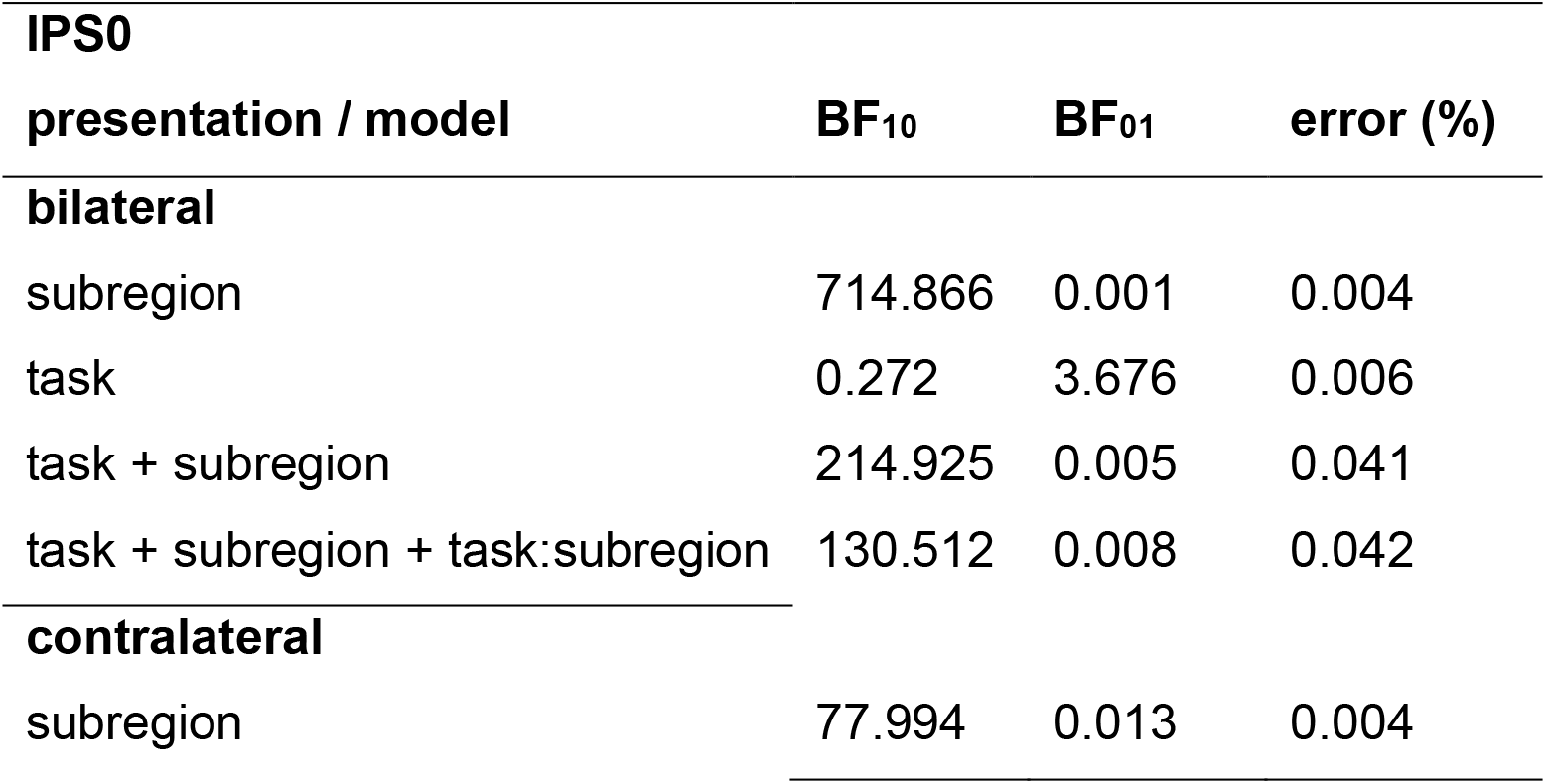
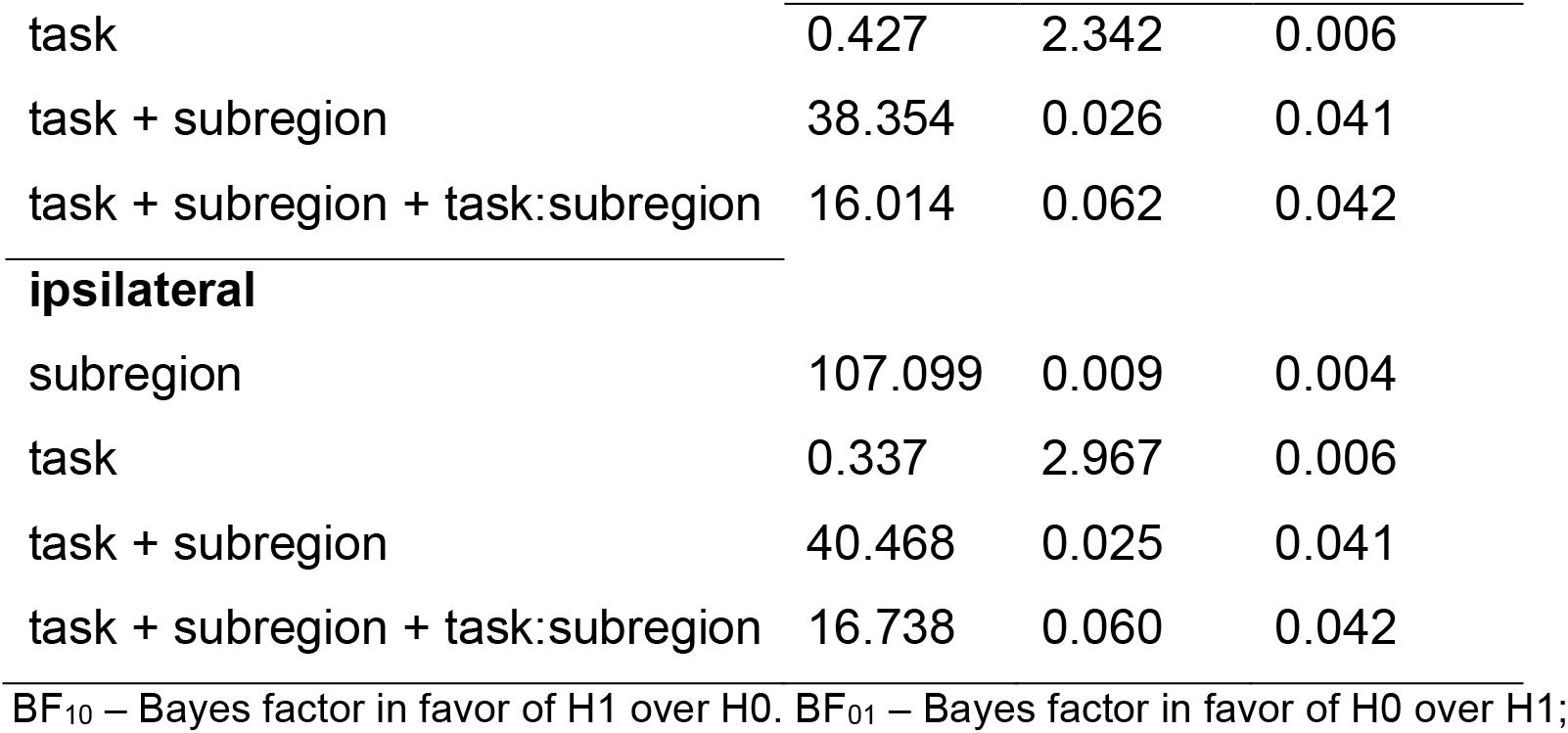
Bayesian evidence for illusory responses in IPS0.

| **IPS0** |  |  |  |  |
| --- | --- | --- | --- | --- |
| **presentation / model** |  | **BF<sub>10</sub>** | **BF<sub>01</sub>** | **error (%)** |
| **bilateral** |  |  |  |  |
| subregion |  | 714.866 | 0.001 | 0.004 |
| task |  | 0.272 | 3.676 | 0.006 |
| task + subregion |  | 214.925 | 0.005 | 0.041 |
| task + subregion + task:subregion |  | 130.512 | 0.008 | 0.042 |
| **contralateral** |  |  |  |  |
| subregion |  | 77.994 | 0.013 | 0.004 |

|  |  |  |  |
| --- | --- | --- | --- |
| task | 0.427 | 2.342 | 0.006 |
| task + subregion | 38.354 | 0.026 | 0.041 |
| task + subregion + task:subregion | 16.014 | 0.062 | 0.042 |
| **ipsilateral** |  |  |  |
| subregion | 107.099 | 0.009 | 0.004 |
| task | 0.337 | 2.967 | 0.006 |
| task + subregion | 40.468 | 0.025 | 0.041 |
| task + subregion + task:subregion | 16.738 | 0.060 | 0.042 |

BF<sub>10</sub> – Bayes factor in favor of H1 over H0. BF<sub>01</sub> – Bayes factor in favor of H0 over H1;

**Supplementary Table 4.**
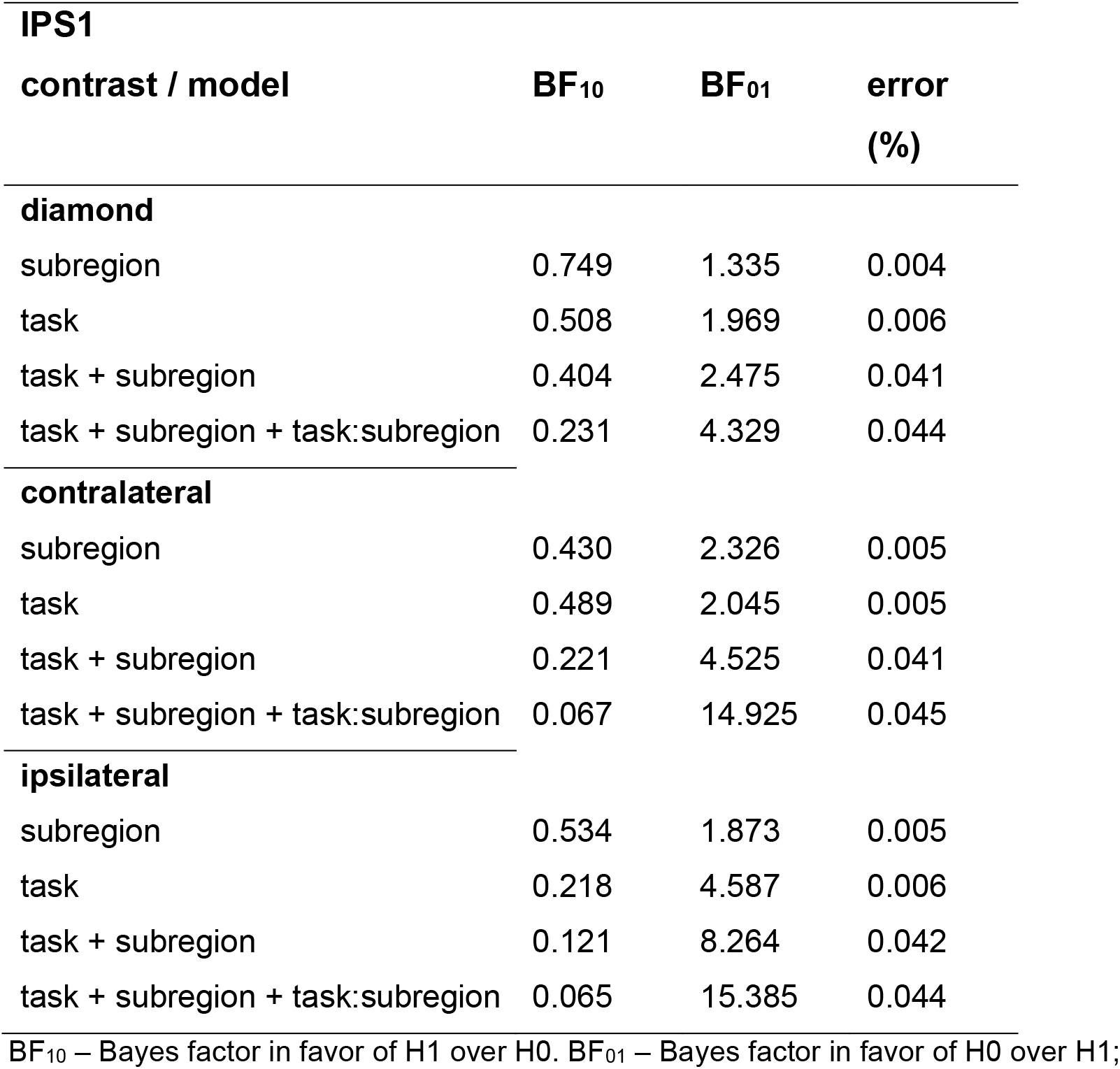
Bayesian evidence for illusory responses in IPS1.

IPS1
| contrast / model | BF<sub>10</sub> | BF<sub>01</sub> | error (%) |
| --- | --- | --- | --- |
| diamond |  |  |  |
| subregion | 0.749 | 1.335 | 0.004 |
| task | 0.508 | 1.969 | 0.006 |
| task + subregion | 0.404 | 2.475 | 0.041 |
| task + subregion + task:subregion | 0.231 | 4.329 | 0.044 |
| contralateral |  |  |  |
| subregion | 0.430 | 2.326 | 0.005 |
| task | 0.489 | 2.045 | 0.005 |
| task + subregion | 0.221 | 4.525 | 0.041 |
| task + subregion + task:subregion | 0.067 | 14.925 | 0.045 |
| ipsilateral |  |  |  |
| subregion | 0.534 | 1.873 | 0.005 |
| task | 0.218 | 4.587 | 0.006 |
| task + subregion | 0.121 | 8.264 | 0.042 |
| task + subregion + task:subregion | 0.065 | 15.385 | 0.044 |

BF<sub>10</sub> – Bayes factor in favor of H1 over H0. BF<sub>01</sub> – Bayes factor in favor of H0 over H1;

**Supplementary Table 5.**
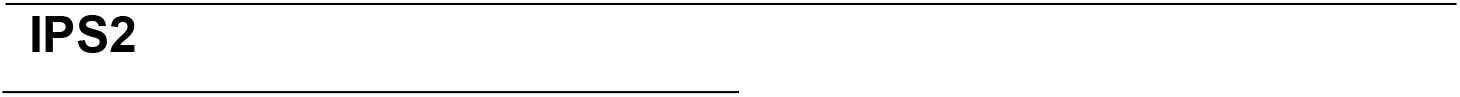
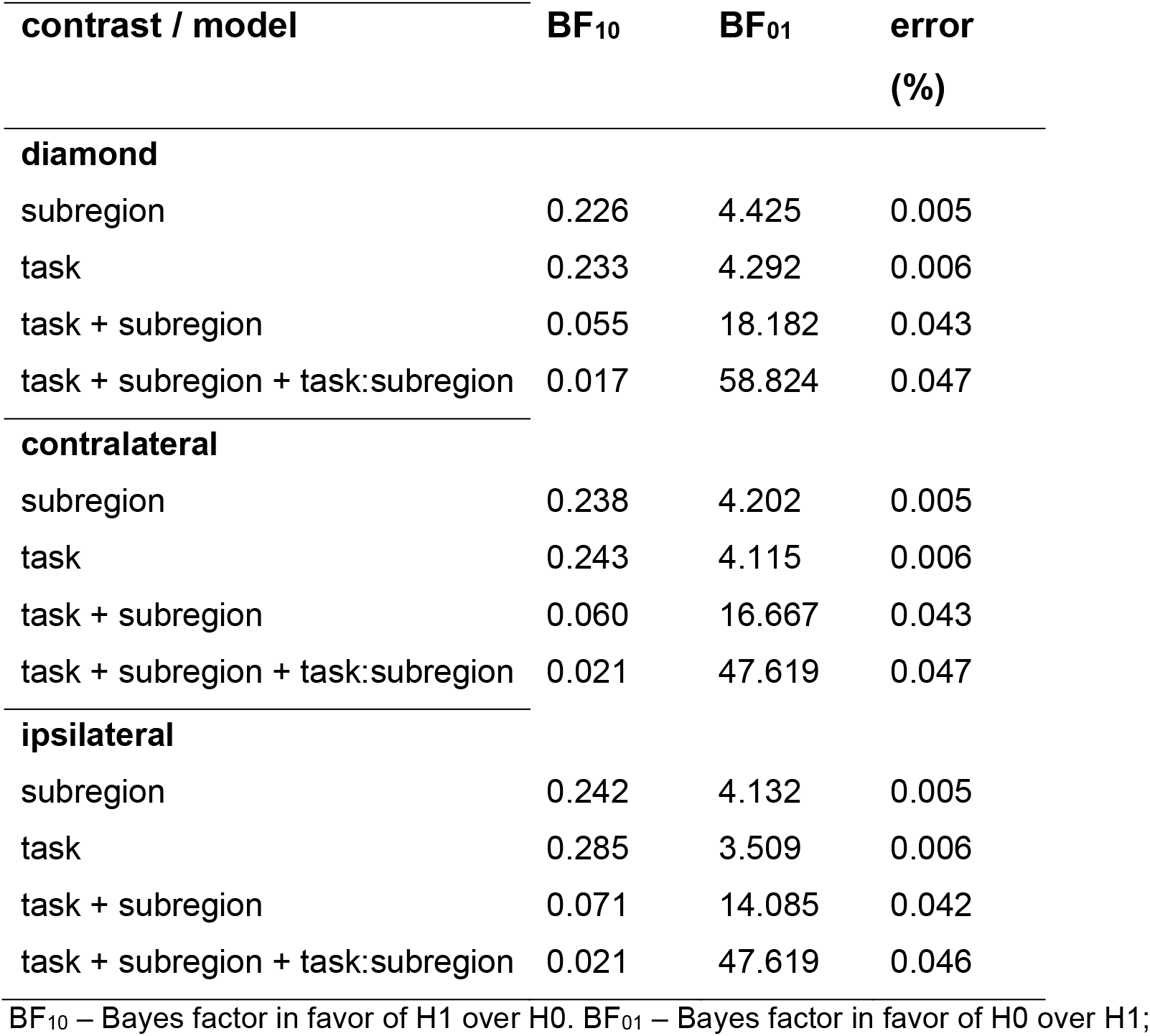
Bayesian evidence for illusory responses in IPS2.

IPS2
| **contrast / model** | **BF<sub>10</sub>** | **BF<sub>01</sub>** | **error (%)** |
| --- | --- | --- | --- |
| **diamond** |  |  |  |
| subregion | 0.226 | 4.425 | 0.005 |
| task | 0.233 | 4.292 | 0.006 |
| task + subregion | 0.055 | 18.182 | 0.043 |
| task + subregion + task:subregion | 0.017 | 58.824 | 0.047 |
| **contralateral** |  |  |  |
| subregion | 0.238 | 4.202 | 0.005 |
| task | 0.243 | 4.115 | 0.006 |
| task + subregion | 0.060 | 16.667 | 0.043 |
| task + subregion + task:subregion | 0.021 | 47.619 | 0.047 |
| **ipsilateral** |  |  |  |
| subregion | 0.242 | 4.132 | 0.005 |
| task | 0.285 | 3.509 | 0.006 |
| task + subregion | 0.071 | 14.085 | 0.042 |
| task + subregion + task:subregion | 0.021 | 47.619 | 0.046 |

BF<sub>10</sub> – Bayes factor in favor of H1 over H0. BF<sub>01</sub> – Bayes factor in favor of H0 over H1;

**Supplementary Table 6.**
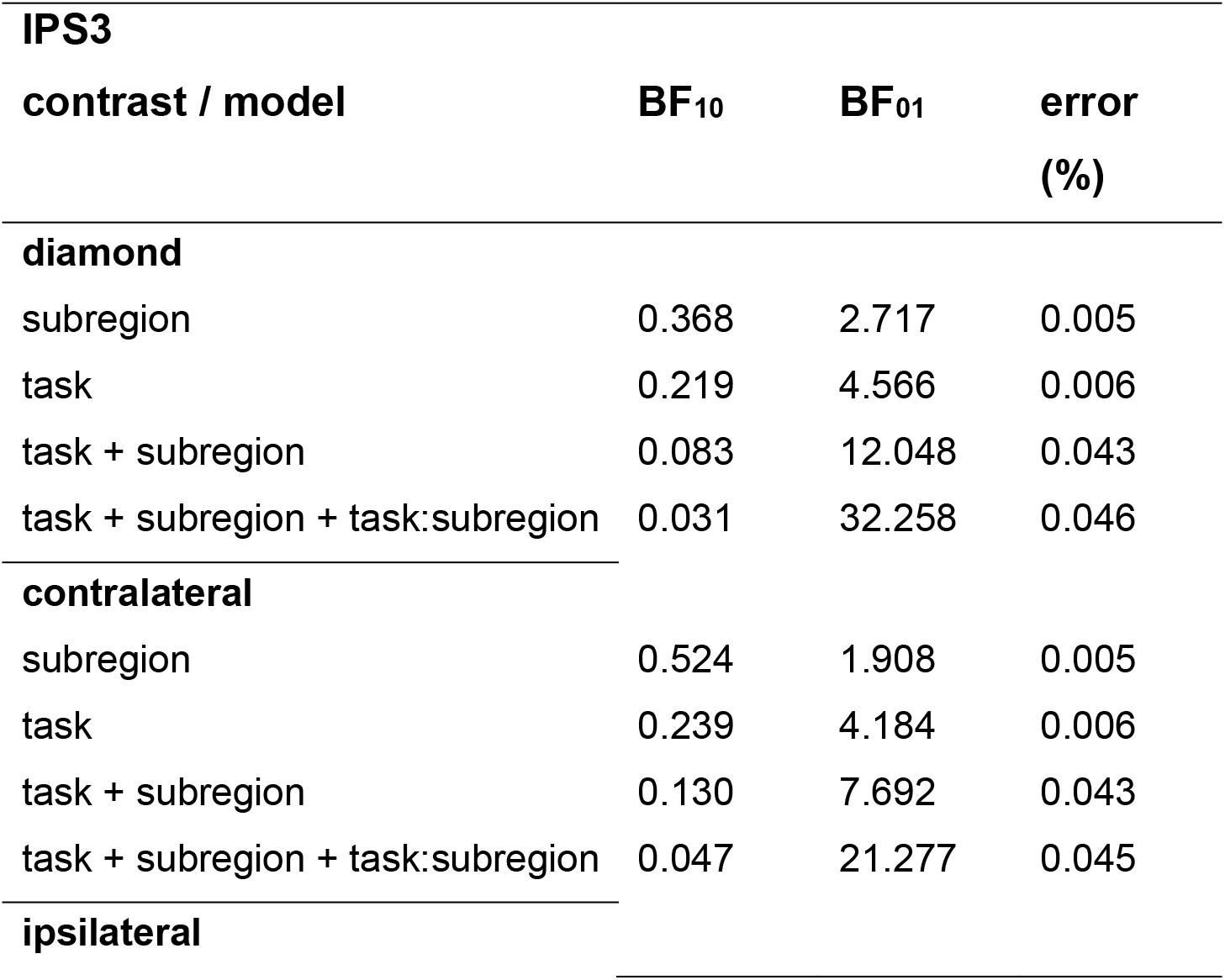
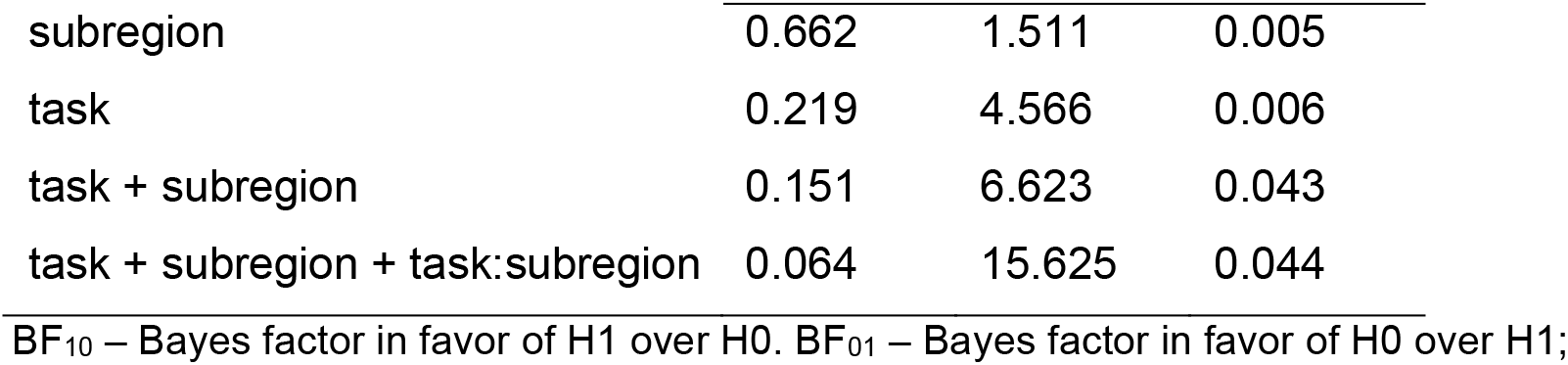
Bayesian evidence for illusory responses in IPS3.

| **IPS3** |  |  |  |
| --- | --- | --- | --- |
| **contrast / model** | **BF<sub>10</sub>** | **BF<sub>01</sub>** | **error (%)** |
| **diamond** |  |  |  |
| subregion | 0.368 | 2.717 | 0.005 |
| task | 0.219 | 4.566 | 0.006 |
| task + subregion | 0.083 | 12.048 | 0.043 |
| task + subregion + task:subregion | 0.031 | 32.258 | 0.046 |
| **contralateral** |  |  |  |
| subregion | 0.524 | 1.908 | 0.005 |
| task | 0.239 | 4.184 | 0.006 |
| task + subregion | 0.130 | 7.692 | 0.043 |
| task + subregion + task:subregion | 0.047 | 21.277 | 0.045 |
| **ipsilateral** |  |  |  |
| subregion | 0.662 | 1.511 | 0.005 |
| task | 0.219 | 4.566 | 0.006 |
| task + subregion | 0.151 | 6.623 | 0.043 |
| task + subregion + task:subregion | 0.064 | 15.625 | 0.044 |

BF<sub>10</sub> – Bayes factor in favor of H1 over H0. BF<sub>01</sub> – Bayes factor in favor of H0 over H1;

**Supplementary Table 7.**
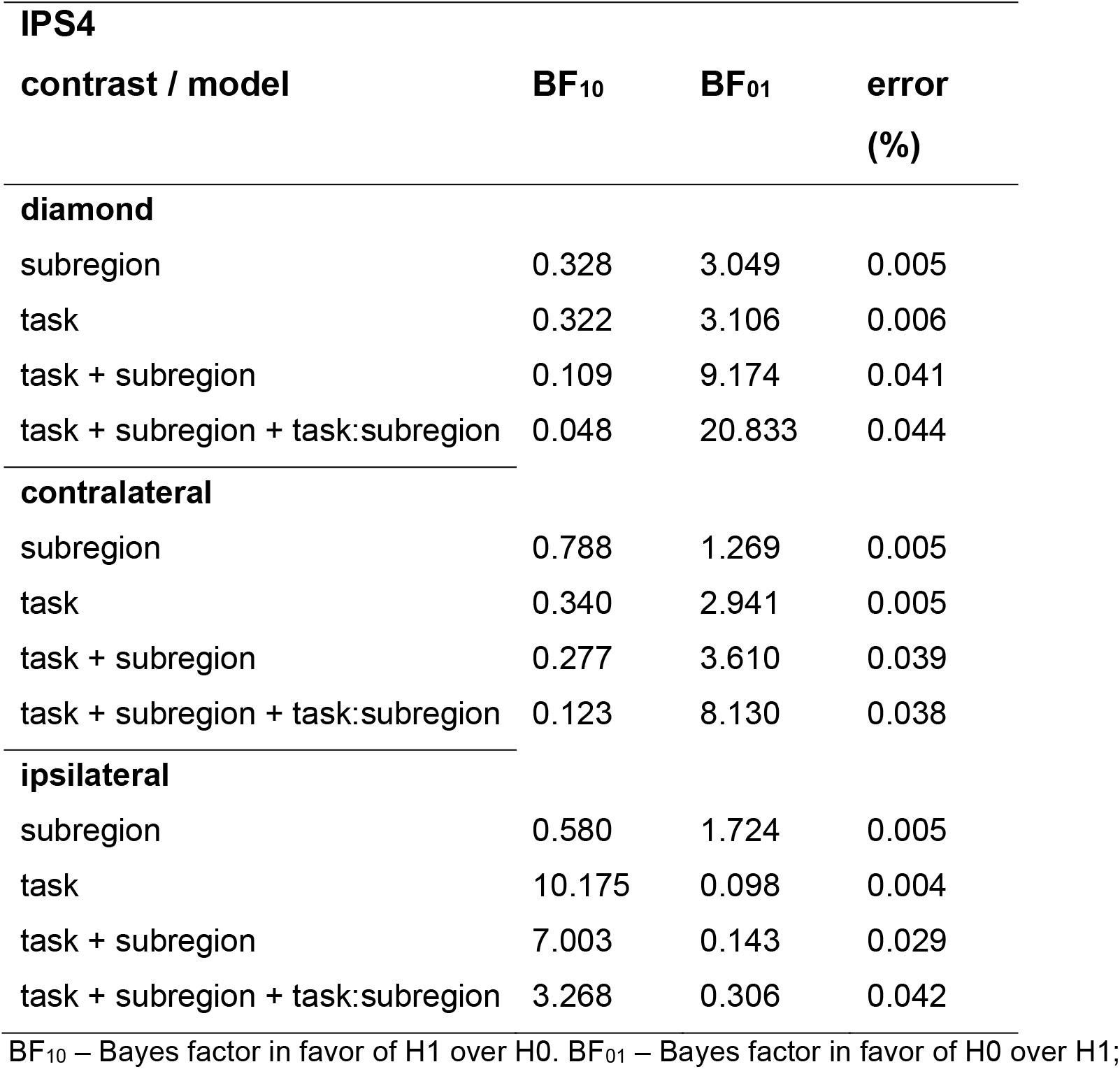
Bayesian evidence for illusory responses in IPS4.

IPS4
| contrast / model | BF<sub>10</sub> | BF<sub>01</sub> | error (%) |
| --- | --- | --- | --- |
| diamond |  |  |  |
| subregion | 0.328 | 3.049 | 0.005 |
| task | 0.322 | 3.106 | 0.006 |
| task + subregion | 0.109 | 9.174 | 0.041 |
| task + subregion + task:subregion | 0.048 | 20.833 | 0.044 |
| contralateral |  |  |  |
| subregion | 0.788 | 1.269 | 0.005 |
| task | 0.340 | 2.941 | 0.005 |
| task + subregion | 0.277 | 3.610 | 0.039 |
| task + subregion + task:subregion | 0.123 | 8.130 | 0.038 |
| ipsilateral |  |  |  |
| subregion | 0.580 | 1.724 | 0.005 |
| task | 10.175 | 0.098 | 0.004 |
| task + subregion | 7.003 | 0.143 | 0.029 |
| task + subregion + task:subregion | 3.268 | 0.306 | 0.042 |

BF<sub>10</sub> – Bayes factor in favor of H1 over H0. BF<sub>01</sub> – Bayes factor in favor of H0 over H1;

